# Shifts in high-light acclimation strategies associated with molecular evolutionary rates in duckweeds

**DOI:** 10.64898/2026.09.07.749874

**Authors:** Yawako W. Kawaguchi, Masaru Kono, Minako Isoda, Kensuke Kawade, Hiromitsu Tabeta, Ryosuke Sasaki, Akira Oikawa, Masami Y. Hirai, Evelina Josephine, Levina Valerie, Natsu Katayama

## Abstract

Molecular evolutionary rates vary widely among lineages, yet the biological processes generating this variation remain poorly understood. In plants, which lack an early-segregated germline, growth- and environment-dependent physiology could itself influence mutation accumulation and, ultimately, substitution rates. Here we examine this possibility in duckweeds (Araceae, Lemnoideae), whose closely related lineages differ severalfold in sequence divergence. Nonsynonymous and synonymous divergence scaled proportionally across all six lineages examined, indicating that this heterogeneity does not reflect differences in selective constraint and instead points to variation in mutation rate. Lineages without prominent anthocyanin accumulation occupied lower latitudes and, under high light, sustained higher relative growth rates and higher photosynthetic efficiency. Comparing the early-diverging, anthocyanin-accumulating *Spirodela polyrhiza* with the late-diverging, anthocyanin-free *Wolffia australiana* revealed contrasting acclimation strategies: *S. polyrhiza* preferentially induced photoprotective pathways, including anthocyanin biosynthesis and non-photochemical quenching, whereas *W. australiana* instead expanded mitochondrial oxidative phosphorylation together with the cytosolic translational machinery, and broadly remodeled primary metabolism. Duckweed lineages therefore differ in how they balance photoprotection against photochemical energy use and downstream metabolic capacity, a divergence that may be associated with their lineage-specific molecular evolutionary rates.

## Introduction

Rates of molecular evolution vary widely among lineages, and understanding the causes of this heterogeneity remains a central issue in evolutionary biology. According to the Neutral Theory and the Nearly Neutral Theory, substitution rates are shaped by the rate at which mutations arise and by their probabilities of fixation, which are influenced by selection and effective population size. The variations in molecular evolutionary rates among lineages is therefore expected to reflect differences in mutation rate, selective constraint, population-genetic environment, or their interactions (Kimura 1983, Ohta 1992). However, despite this strong theoretical framework, experimental studies that explain molecular evolutionary rate variation in natural lineages by linking it to ecological or physiological traits remain scarce, particularly in plants (Smith and Donoghue, 2008; Lanfear et al., 2010).

In flowering plants, substantial variation in molecular evolutionary rates has been reported across lineages and has been associated with differences in life history (Smith & Donoghue 2008, Smith 2025), specialized trophic strategy as parasitism or carnivory (Müller et al 2004, Ibarra-Laclette et al 2011, Bromham et al 2013) and habitat environment (Katayama et al 2022, Quian et al 2024). However, despite these associations, a mechanistic understanding of how life-history traits causally influence molecular evolutionary rates is still limited. In particular, although theoretical models predict that variation in mutation rate should contribute to differences in substitution rates, empirical studies examining whether and how life-history traits affect mutation rates are scarce. In addition, reproduction modes of plants do not have an early segregation of germline and imply that somatic cell divisions may contribute to heritable mutations (Lanfear et al 2018). This biological feature suggests that differences in growth rate and cumulative cell division could influence mutation rate. Plants characterized by rapid growth or metabolic turnover may experience increased DNA replication events or oxidative stress, potentially elevating mutation rates and, consequently, accelerating molecular evolutionary rates.

Light availability is particularly relevant in this context because it directly affects both carbon gain and physiological stress in plants. Photosynthesis provides the energy and carbon required for growth (Smith and Stitt, 2007), and photosynthetic capacity and efficiency are therefore closely associated with growth rate, as plant growth depends on carbon assimilation to sustain continuous cell proliferation (Schurr et al., 2006; Smith et al., 2024). Because mutation accumulation is ultimately linked to DNA replication, variation in growth rate may indirectly contribute to differences in molecular evolutionary rates (Lanfear, 2018; Watson et al., 2016). At the same time, when absorbed light exceeds photosynthetic demand, excess of light energy can cause over-reduction of the photosynthetic electron transport chain and impose oxidative stress (Foyer and Shigeoka, 2011; Khorobrykh et al., 2020). Plants must therefore balance the use of light energy for photochemistry with photoprotective processes to avoid photoinhibition (Ruban, 2016; Didaran et al., 2024).

Plants have evolved diverse strategies to manage this trade-off under high-light environments (Zhang et al., 2024). These include enhanced pigment accumulation and non-photochemical quenching (NPQ), as well as increased photorespiration, mitochondrial respiration, and other metabolic sinks that dissipate excess reducing equivalents (Ort and Baker, 2002; Ruban, 2016; Demircan et al., 2024; Zhang et al., 2024). The relative contribution of these processes is likely to vary among lineages and would explain differences in growth performance under high light (Paul and Foyer, 2001; Rosado-Souza et al., 2023). If so, lineage-specific differences in high-light acclimation may also be linked to variation in growth rate and, ultimately, molecular evolutionary rates.

In this study, we focused on the duckweed subfamily (Lemnoideae, Araceae), a group of free-floating monocotyledonous aquatic plants known as the world’s smallest flowering plants, because phylogenetically, duckweeds exhibit exceptionally long branch lengths compared with other members of Araceae (Rothwell et al 2004; Nauheimer et al, 2012). The subfamily comprises five genera—*Spirodela*, *Landoltia*, *Lemna*, *Wolffiella*, and *Wolffia* and molecular phylogenetic analyses indicate that *Spirodela* diverged first, followed by *Landoltia* and then *Lemna*, whereas *Wolffiella* and *Wolffia* form the last-branching clade (Acosta et al., 2021). Along this phylogenetic relationship, variation in substitution rates was observed within the subfamily, with the tendency that later-branching lineages show higher rates (Nauheimer et al, 2012). Duckweeds also show highly simplified body plan consisting of small leaf-like organs called “fronds,” which bear roots or root-like structures on the ventral side (Fig. 1A). These fronds float on the water surface and propagate clonally through repeated budding. Owing to their minute body size (<1–15 mm), duckweeds exhibit exceptionally rapid growth rates among flowering plants (Sree et al., 2015; Ziegler et al., 2015). They are nearly cosmopolitan in distribution, occurring on most continents except in arid regions and polar areas (Landolt 1986; Nauheimer et al 2012). In natural habitats, they grow in open freshwater environments, where they are exposed to either direct sunlight or the shade of surrounding aquatic vegetation. Although interspecific differences in ecophysiological traits have been reported (Burgess et al., 2023; Smith et al., 2024), the ecophysiological evolutionary processes underlying adaptation to diverse light environments remain poorly understood.

**Fig. 1.**
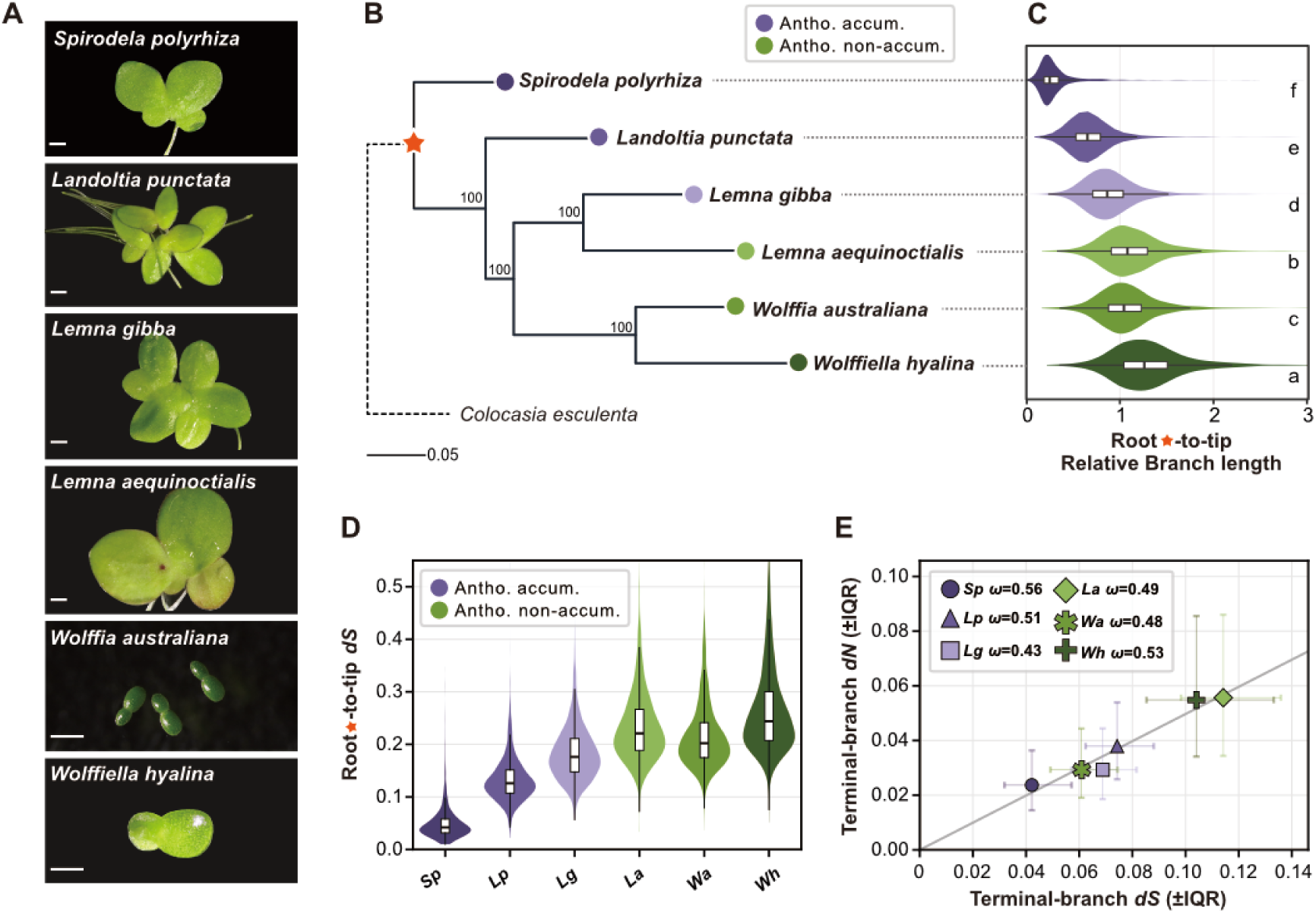
Phylogeny and lineage-specific molecular evolutionary rates in duckweeds. (A) The six Lemnoideae lineages used in this study. Scale bars, 1 mm. (B) Maximum-likelihood species tree inferred from the concatenated codon alignment of 2,819 single-copy orthogroups, with *Colocasia esculenta* as an outgroup. Numbers at nodes are ultrafast bootstrap support values; the outgroup branch (dashed) is not to scale. The orange star marks the root node, defined as the split between *C. esculenta* and the six Lemnoideae species, from which root-to-tip distances in (C) and (D) were measured. Scale bar, 0.05 substitutions per site. (C) Relative molecular evolutionary rate of each species, aligned to the tree in (B). For each gene tree, the root-to-tip distance of a species was divided by that of *C. esculenta* in the same tree. Violins show the distribution across gene trees and boxes give the median and interquartile range. Letters (a–f) denote statistically distinct groups (Friedman test followed by post-hoc pairwise Wilcoxon signed-rank tests with Holm correction; all pairs *p* < 0.05), ordered by decreasing median rate. (D) Root-to-tip synonymous divergence (*d_S_*) per species, estimated under the MG94×REV codon model across 2,474 single-copy orthogroups passing quality control. Violins and boxes as in (C). (E) Median terminal-branch nonsynonymous (*d_N_*) against synonymous (*d_S_*) divergence for each species; error bars span the interquartile range across orthogroups. The grey line is the proportional fit through the origin (constant *d_N_/d_S_*), giving *ω* = 0.50. Species abbreviations: *Sp*, *Spirodela polyrhiza*; *Lp*, *Landoltia punctata*; *Lg*, *Lemna gibba*; *La*, *Lemna aequinoctialis*; *Wa*, *Wolffia australiana*; *Wh*, *Wolffiella hyalina*.

In the present study, we examined ecophysiological responses to light to infer factors that may be associated with lineage-specific molecular evolutionary rate variation in duckweeds. We first compared relative molecular evolutionary rates across six duckweed species and then examined the geographic distributions of a broader set of duckweed species and found that distribution patterns differed between anthocyanin-accumulating and anthocyanin-non-accumulating lineages. We then compared growth rates, photoinhibition, and photosynthetic responses under low-light (LL) and high-light (HL) conditions between anthocyanin-accumulating and anthocyanin-non-accumulating lineages and further analyzed transcriptomic and metabolomic responses to high light in *Spirodela polyrhiza* and *Wolffia australiana*, which are representative of anthocyanin-accumulating and anthocyanin-non-accumulating lineages, respectively. By integrating phylogenomic, physiological, and molecular data, we aimed to clarify how lineage-specific ecological and physiological strategies under high light are associated with molecular evolutionary rate variation in these exceptionally small and rapidly growing plants.

## Results

### Comparison of molecular evolutionary rates

To examine variation in molecular evolutionary rates within duckweeds, we analyzed 2,819 single-copy orthogroups identified from six Lemnoideae species (*Spirodela polyrhiza* (L.) Schleid., *Landoltia punctata* (G.Mey.) Les & D.J.Crawford, *Lemna aequinoctialis* Welw., *Lemna gibba* L., *Wolffia australiana* (Benth.) Hartog & Plas, and *Wolffiella hyalina* (Delile) Monod; Fig. 1A) and *Colocasia esculenta* (L.) Schott as an outgroup. Using codon alignments of these orthogroups, we inferred a maximum-likelihood species tree within the subfamily (Fig. 1B). Consistent with previous studies, *Spirodela* diverged first, followed by *Landoltia*, then the two *Lemna* species, whereas *Wolffiella* and *Wolffia* formed the last-branching clade. To compare relative molecular evolutionary rates among lineages, we calculated root-to-tip branch lengths for each species across gene trees and normalized them by the corresponding branch length of *C. esculenta* in the same tree. Relative branch lengths differed significantly among all six duckweed species (Fig. 1C; all pairs *p* < 0.05), indicating substantial heterogeneity in sequence divergence across Lemnoideae lineages. The difference was most pronounced between *S. polyrhiza* and *Wolffiella hyalina*, whose median relative branch lengths differed approximately five-fold (0.250 and 1.260, respectively).

To test whether this rate heterogeneity reflects lineage-specific selection or a largely neutral process, we estimated synonymous and nonsynonymous substitutions (*d_S_* and *d_N_*, respectively) across 2,474 single-copy orthologues passing quality control. Root-to-tip synonymous divergence (*d_S_*) from the duckweed common ancestor, which controls for evolutionary time, varied approximately six-fold among lineages (Fig. 1D), consistent with the branch-length variation in Fig. 1C. To ask whether this reflected differences in selection, we compared *d_S_* and *d_N_* on each species’ terminal branch. Median *d_N_* scaled proportionally with median *d_S_* (slope through the origin, *d_N_/d_S_* (*ω*) = 0.50), and all six lineages fell on this common line (Fig. 1E). Per-lineage ω values were tightly clustered, ranging only from 0.43 in *L. gibba* to 0.56 in *S. polyrhiza* — a 1.3-fold spread with broadly overlapping interquartile ranges, against the approximately five-fold difference in divergence itself. Thus the several-fold variation in evolutionary rate among duckweed lineages is not accompanied by a difference in selection pressure, indicating that it is largely neutral and consistent with variation in the underlying mutation rate rather than in selection.

### Geographic distribution of anthocyanin-accumulating and anthocyanin-non-accumulating duckweeds

Previous studies have shown that anthocyanin production varies among duckweed lineages. (Les et al., 1997). Members of the genera *Spirodela* and *Landoltia* produce anthocyanins, whereas anthocyanin production has not been observed in members of *Wolffiella* and *Wolffia*. In addition, species of *Lemna* are divided into anthocyanin-accumulating and anthocyanin-non-accumulating clades based on the molecular phylogeny and flavonoid profile (Bog et al. 2020a, 2020b, Les et al., 1997). Because anthocyanins can function in protection against oxidative stress, particularly under excessive light conditions (Gould, 2004; Landi et al., 2015), lineage-specific differences in anthocyanin production may reflect ecological characteristics among duckweed lineages. To explore potential ecological differentiation among lineages, we mapped global occurrence records of duckweed species obtained from GBIF (Fig. 2; GBIF.org,19 September 2024). After species-level filtering, coordinate cleaning, duplicate removal, and spatial thinning to one occurrence per species within each 0.5-degree grid cell, the dataset contained 18,227 occurrence records from 29 species (Data S1, S2). Compared with lineages that do not produce anthocyanins, anthocyanin-accumulating lineages were distributed at higher latitudes and in more inland regions (median absolute latitude, 48.3 degrees for anthocyanin-accumulating species and 37.5 degrees for anthocyanin-non-accumulating species; Figs. 2A, 2B). A similar pattern was observed within the genus *Lemna*, whereanthocyanin-accumulating species exhibited a broader distribution extending to higher latitudes and inland regions than anthocyanin-non-accumulating species (Fig. S1). These results suggest that anthocyanin-accumulating duckweed lineages may occupy a wider geographic range, including higher-latitude and inland environments, whereas anthocyanin-non-accumulating lineages may be relatively more associated with lower-latitude regions.

**Fig. 2.**
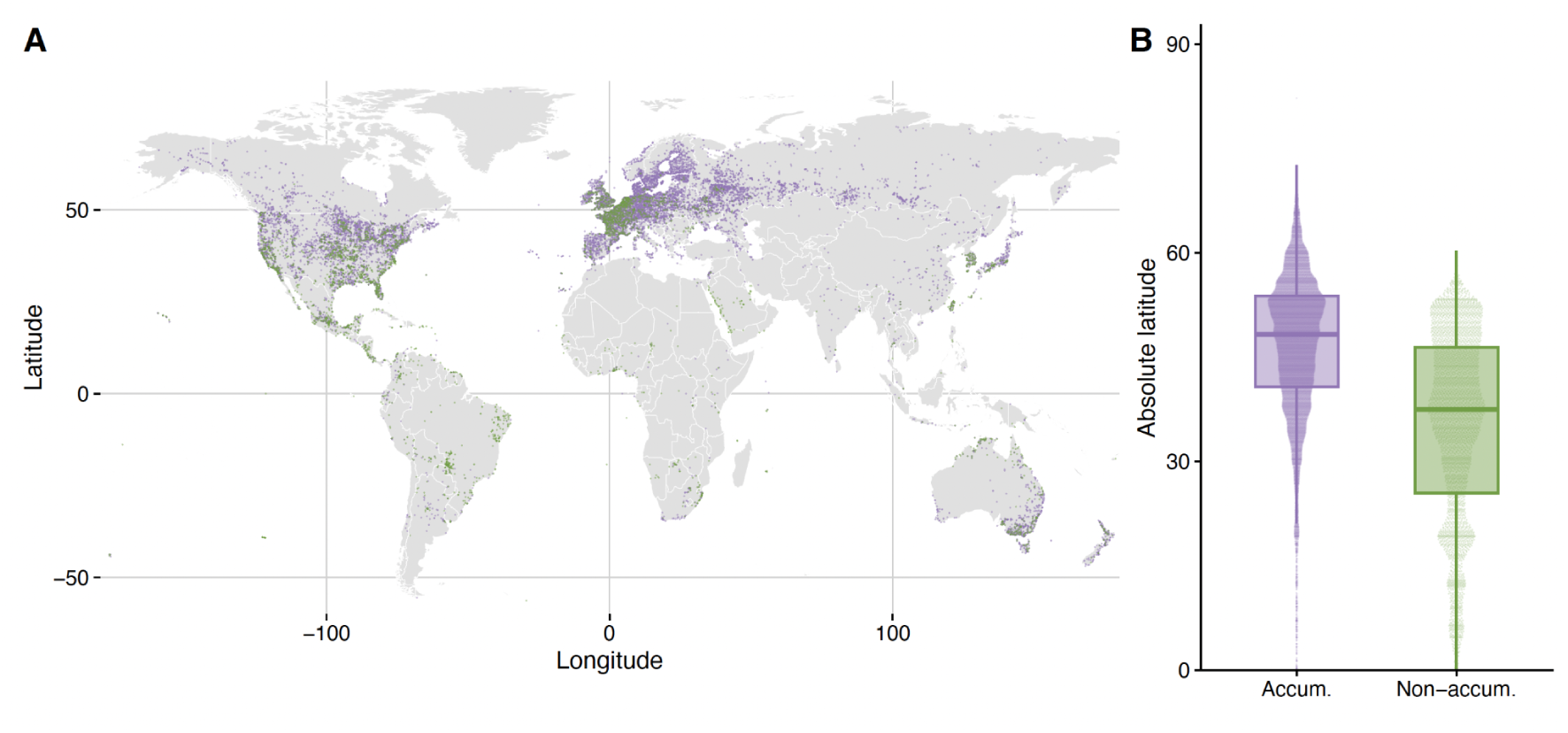
Global distribution of and latitudinal patterns of anthocyanin-accumulating and anthocyanin-non-accumulating duckweed species. (A) Global distribution of duckweed occurrence records obtained from GBIF. Records were filtered to one occurrence per species within each 0.5-degree grid cell. Purple points indicate anthocyanin-accumulating species and green points indicate anthocyanin-non-accumulating species, based on Les et al. (1997). (B) Absolute latitude distributions of anthocyanin-accumulating and anthocyanin-non-accumulating species calculated from the same thinned occurrence dataset. Boxplots indicate median and interquartile range, and overlaid points indicate thinned occurrence records.

### Growth rate comparisons among lineages under different light conditions

Because anthocyanins are associated with protection against excess light, the occurrence of anthocyanin-non-accumulating lineages in lower-latitude regions raises the question of whether these lineages could cope with potentially stronger light environments. To address this question, we first compared growth rates under LL and HL conditions and tested whether anthocyanin-non-accumulating lineages can grow under high light conditions.

We measured frond area at approximately 24-h intervals for seven days in six representative duckweed accessions grown under LL (approximately 50–100 µmol m⁻² s⁻¹ PFD) and HL (approximately 800–1000 µmol m⁻² s⁻¹ PFD) conditions (Data S3). To evaluate the growth rates, we fitted both exponential and logistic growth-curve models to each well and compared model fit using AIC (Fig. S2, Data S4). The AIC-based model comparison indicated that some wells were better described by logistic rather than purely exponential growth when later time points were included, consistent with approach to carrying capacity. We therefore used the Day 0 to Day 4 interval, which captured a common early growth phase across lineages and light conditions, to calculate 4 Day relative growth rate (RGR) for the main comparison (Fig. 3A; Data S5). All six accessions showed higher RGR under HL than under LL, but the magnitude of this increase differed among lineages. The HL/LL RGR ratio ranged from 1.29 in *Landoltia punctata* to 1.85 in *Wolffiella hyalina*, with relatively large increases in the anthocyanin-non-accumulating lineages *Wolffiella hyalina* and *W. australiana* (Fig. 3A; Data S5).

**Fig. 3.**
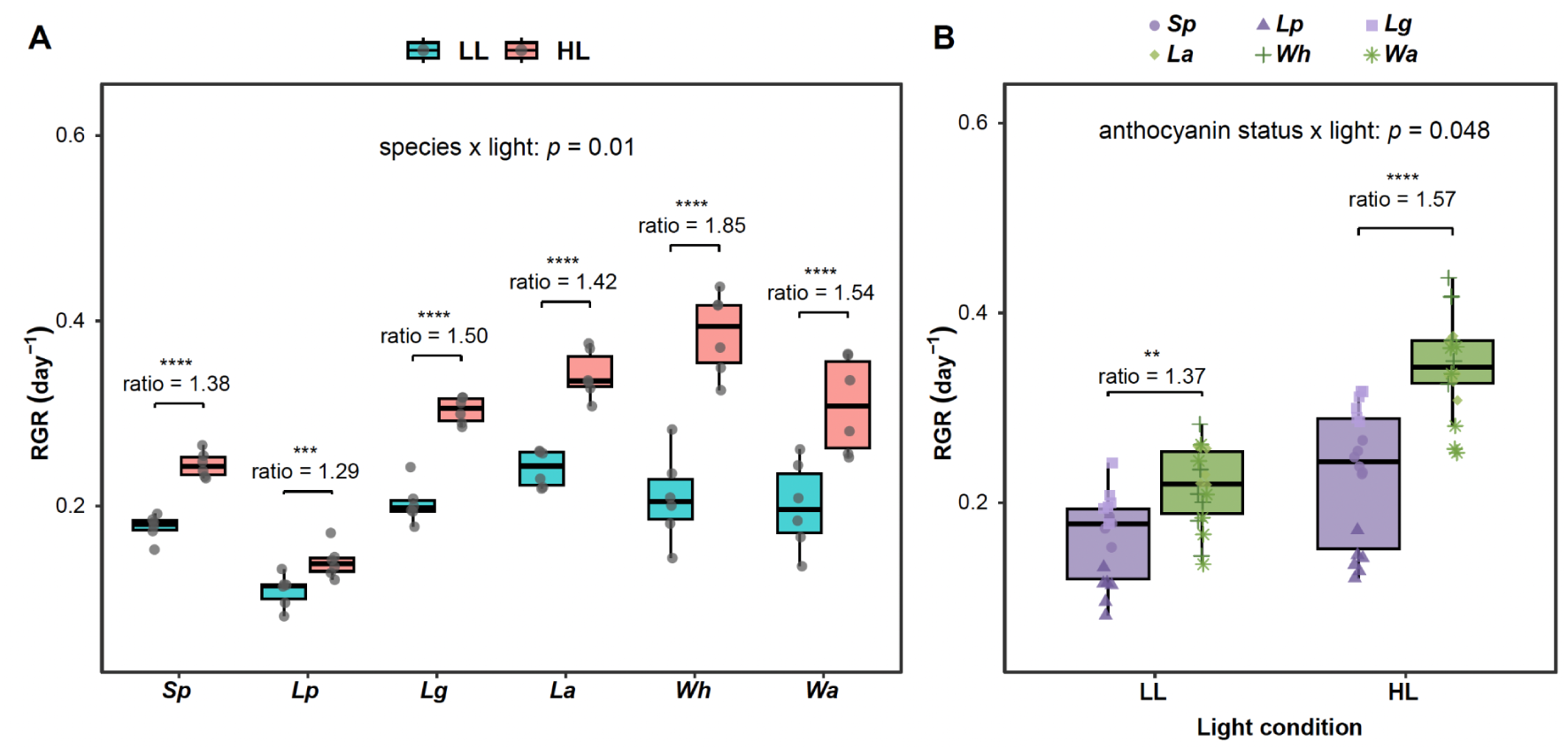
Relative growth rate (RGR) under low-light (LL) and high-light (HL) conditions. (A) RGR of six duckweed species under LL (approximately 50–100 µmol m⁻² s⁻¹) and HL (approximately 800–1000 µmol m⁻² s⁻¹). Points represent biological replicates and boxplots indicate median and interquartile range. (B) Estimated RGR for anthocyanin-accumulating and anthocyanin-non-accumulating lineages under LL and HL conditions. anthocyanin-accumulating lineages are shown in purple and anthocyanin-non-accumulating lineages are shown in green. Estimated marginal means were calculated from a Gamma generalized linear mixed-effects model including light condition, anthocyanin status, and their interaction as fixed effects, with species treated as a random effect. Error bars represent standard errors.

Consistent with these observations, RGR was significantly affected by anthocyanin production status, light intensity, and their interaction in a Gamma generalized linear mixed-effects model (anthocyanin status: *χ²* = 17.40, *p* < 0.001; light intensity: *χ²* = 130.97, *p* < 0.001; anthocyanin status x light intensity: *χ²* = 3.91, *p* = 0.048; Data S5). Estimated marginal means showed that anthocyanin-non-accumulating lineages had higher RGR than anthocyanin-accumulating lineages under both LL and HL conditions, with a stronger difference under HL. Under LL, the estimated RGR was 0.158 day^-1 for anthocyanin-accumulating lineages and 0.216 day⁻¹ for anthocyanin-non-accumulating lineages (non-antho/antho ratio = 1.37, *p* = 0.001). Under HL, the estimated RGR was 0.220 day^-1 for anthocyanin-accumulating lineages and 0.344 day^-1 for anthocyanin-non-accumulating lineages (non-antho/antho ratio = 1.57, *p* < 0.001) (Fig. 3B; Data S5). These results indicate that anthocyanin-non-accumulating lineages maintain higher growth rates under both light environments, and that this difference is accentuated under high light.

### Lineage-specific photoinhibition and recovery under high light

Next, to evaluate tolerance to high light across duckweed lineages, we measured the maximum quantum yield of PSII photochemistry (F_v_/F_m_) as an indicator of photoinhibition in the same six representative lineages: *S. polyrhiza*, *Landoltia punctata*, *L. gibba*, *L. aequinoctialis*, *Wolffiella hyalina*, and *W. australiana* (Fig. 4A, Fig. S3, Data S6-1). Plants were precultured under LL for 3 days and then transferred to HL. For the main analysis, we focused on F_v_/F_m_ values at Day 0, Day 1, and Day 4 from the transfer to HL, capturing the initial state, acute photoinhibition, and subsequent recovery phase.

**Fig. 4.**
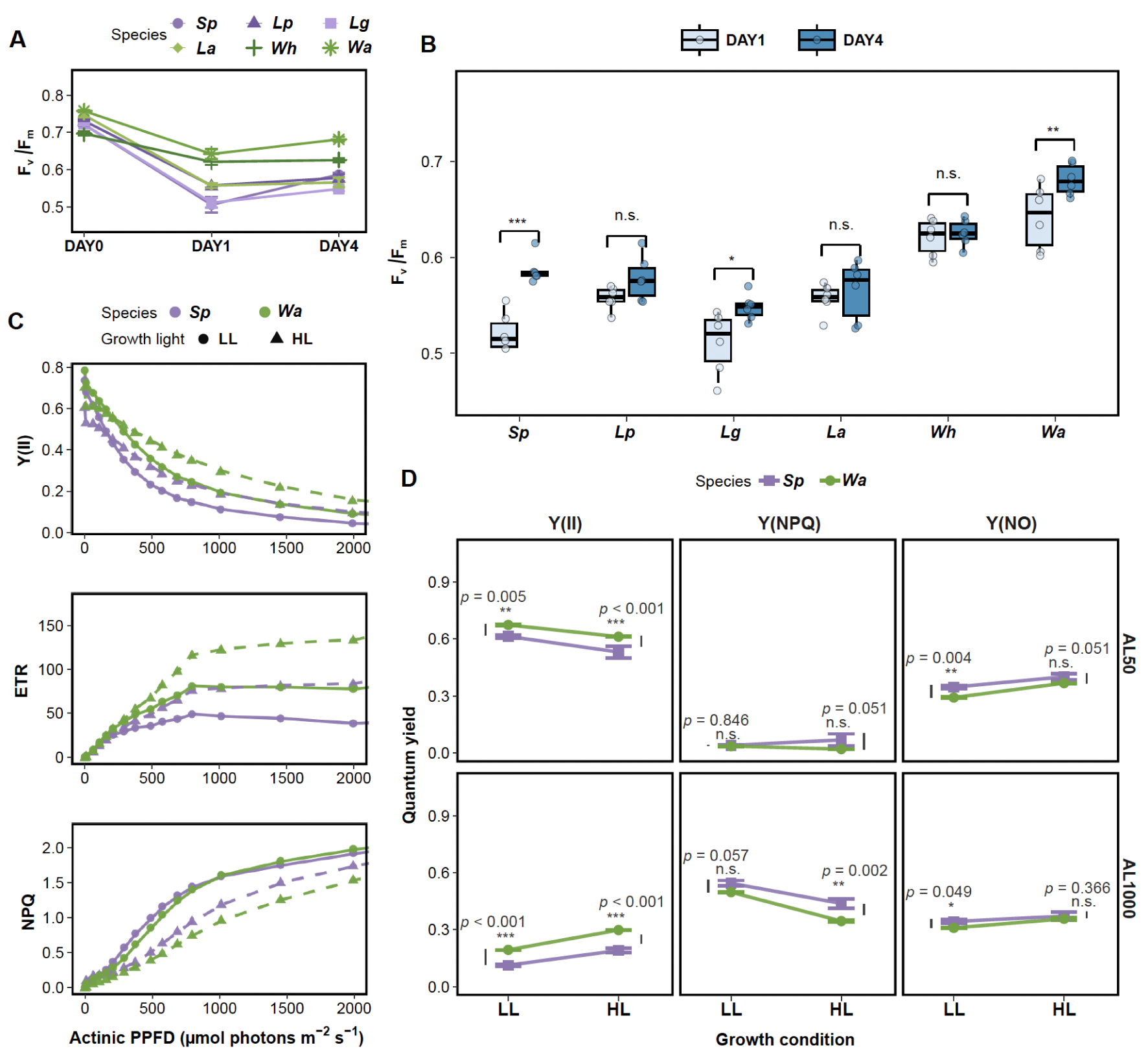
Chlorophyll fluorescence responses to high light. (A) Time course of maximum quantum yield of PSII photochemistry (Fv/Fm) after transfer to high light. Points and error bars indicate mean +/- SE for six representative duckweed accessions at Day 0, Day 1, and Day 4. (B) Within-lineage comparison of Fv/Fm between Day 1 and Day 4 after transfer to high light. Boxes show the interquartile range and median, points show individual measurements, and brackets indicate Tukey-adjusted Day 1 versus Day 4 contrasts within each lineage. \**p* < 0.05, \*\**p* < 0.01, \*\*\**p* < 0.001; n.s., not significant. (C) Light-response curves of Y(II), ETR, and NPQ in *Spirodela polyrhiza* and *Wolffia australiana* grown under LL or HL conditions. Curves show photosynthetic responses across increasing actinic light intensity. Colors indicate species (*S. polyrhiza*, purple; *W. australiana*, green), and point shapes and line types indicate growth-light condition (LL, circles, solid-line; HL, triangles, dashed-line). (D) Energy partitioning at two representative AL intensities. Y(II), Y(NPQ), and Y(NO) indicate PSII photochemistry, regulated non-photochemical energy dissipation, and non-regulated energy loss, respectively. Bars represent mean +/- SE, and points indicate individual replicates. Brackets indicate pairwise comparisons between species within each growth-light and measurement-light combination; p values are shown with significance levels (* *p* < 0.05, ** *p* < 0.01, *** *p* < 0.001; n.s., not significant).

Initial Fv/Fm values before HL exposure were high across lineages but differed modestly among lineages (Fig. 4A, Fig. S3, Data S6-2). *W. australiana* showed the highest mean Fv/Fm at Day 0 (0.758 +/- 0.003 SE), whereas *Wolffiella hyalina* showed the lowest Day 0 value (0.697 +/- 0.006 SE). After one day under HL, Fv/Fm decreased in all accessions, indicating photoinhibition or down-regulation of PSII. However, the magnitude of this decrease differed strongly among lineages. The anthocyanin-non-accumulating lineages *W. australiana* and *Wolffiella hyalina* retained the highest Fv/Fm values at Day 1 (0.642 +/- 0.014 and 0.621 +/- 0.008, respectively), whereas *S. polyrhiza* and *L. gibba* showed lower Day 1 values (0.505 +/- 0.021 and 0.511 +/- 0.013, respectively).

By Day 4, F_v_/F_m_ had partially recovered in several accessions, but recovery patterns remained lineage-specific. *W. australiana* retained the highest mean F_v_/F_m_ at Day 4 (0.681 +/- 0.007), followed by *Wolffiella hyalina* (0.626 +/- 0.006), whereas *L. gibba* showed the lowest Day 4 value among the six accessions (0.548 +/- 0.005) (Data S6-2). Within each lineage, we compared Fv/Fm between Day 1 and Day 4 to evaluate recovery after the initial decline under HL. Fv/Fm increased significantly from Day 1 to Day 4 in *S. polyrhiza*, *L. gibba*, and *W. australiana*, but not in *Landoltia punctata*, *L. aequinoctialis*, or *Wolffiella hyalina* (Fig. 4B, Data S6-3). Consistent with these patterns, Fv/Fm was significantly affected by lineage, time, and their interaction under HL conditions (lineage: *F* = 48.76, *p* < 0.001; time: *F* = 569.02, *p* < 0.001; lineage x time: *F* = 12.99, *p* < 0.001; Data S6-4). Tukey-adjusted comparisons among lineages within each time point further supported lineage-specific differences in photoinhibition and recovery (Fig. S3; Data S6-5). These results demonstrate that duckweed lineages differ not only in the magnitude of acute high-light-induced photoinhibition but also in their subsequent recovery dynamics, with *W. australiana* maintaining the highest PSII maximum efficiency after HL exposure. In the following analyses, we focused on *W. australiana* and compared it with *S. polyrhiza* to investigate the physiological and molecular basis of this difference in high-light tolerance.

### Photosynthetic light-response curves and PSII energy partitioning

To further examine physiological differences between *S. polyrhiza* and *W. australiana* under high light, we compared PAM light-response curves after plants had been grown for four days under LL or HL conditions (Fig. 4C). Y(II), which represents the fraction of absorbed light energy used for PSII photochemistry, decreased as actinic light (AL) increased in both species and growth-light conditions. Across the examined AL range, *W. australiana* maintained higher Y(II) than *S. polyrhiza*. In both species, LL-grown plants showed higher Y(II) at low AL, whereas HL-grown plants tended to maintain higher Y(II) under stronger AL. ETR, which represents electron transport through PSII, increased with AL and differed among species and growth-light conditions. HL-grown *W. australiana* showed the highest ETR, LL-grown *S. polyrhiza* showed the lowest ETR, and HL-grown *S. polyrhiza* and LL-grown *W. australiana* showed intermediate values. NPQ, which represents regulated dissipation of excess light energy as heat, increased as AL became stronger in both species and growth-light conditions. At high AL, *S. polyrhiza* tended to show higher NPQ than *W. australiana*, particularly after HL growth, whereas *W. australiana* maintained lower NPQ while retaining higher Y(II). Taken together, these light-response curves indicate that *W. australiana* maintained higher PSII photochemical performance than *S. polyrhiza* across a broad range of AL intensities, particularly after HL acclimation.

We next compared quantum-yield partitioning at two representative actinic light intensities: AL50 (61.885 µmol photons m⁻² s⁻¹) and AL1000 (1012.500 µmol photons m⁻² s⁻¹) (Fig. 4D, Data S7-1). Species comparisons within each growth-light and actinic-light condition showed that *W. australiana* generally maintained higher Y(II) than *S. polyrhiza*. This difference was significant at AL50 in both LL-grown plants (*p* = 0.005) and HL-grown plants (*p* < 0.001), and at AL1000 in both LL- and HL-grown plants (both p < 0.001; Fig. 4D, Data S7-2). For Y(NPQ), differences between species were detected at AL1000, particularly in HL-grown plants, where *S. polyrhiza* showed higher Y(NPQ) than *W. australiana* (*p* = 0.002; Fig. 4D, Data S7-2). These patterns suggest that *S. polyrhiza* depends more strongly on regulated dissipation of excess light energy, whereas *W. australiana* maintains a larger fraction of absorbed light energy in PSII photochemistry under high-light-acclimated conditions. Species x growth-light interaction tests within each actinic-light condition did not detect significant interactions for Y(II), Y(NPQ), or Y(NO) at either AL50 or AL1000 (all p > 0.05; Data S7-3). Together, these results suggest that both species acclimate to growth-light environments, but *W. australiana* maintains higher PSII photochemical yield, whereas *S. polyrhiza* tends to allocate more absorbed energy to regulate non-photochemical dissipation at high light conditions.

### Lineage-specific gene expression in response to high light

To investigate the molecular basis of the contrasting high-light responses in *S. polyrhiza* and *W. australiana*, we performed RNA-Seq analyses under low light (LL), 3 hours after transfer to high light (HL_3h), and 4 days after transfer to high light (HL_4d). Because baseline expression levels differed substantially between the species, we summarized expression at the orthogroup level and compared treatment-dependent expression changes within each species (Data S0-2).

To characterize how *S. polyrhiza* and *W. australiana* differ in their transcriptional response to high light, we classified the high-light response trajectory of each of the 10,916 orthogroups expressed in both species across LL, HL_3h, and HL_4d. Each of the two consecutive transitions (LL→HL_3h and HL_3h→HL_4d) was scored as up (U), flat (F), or down (D) from within-species *limma* contrasts (BH-adjusted *p* < 0.05 and |log2 fold change| ≥ 0.5), yielding nine possible trajectories. The two species deployed temporally distinct response strategies (Fig. 5A, Data S8-1). *S. polyrhiza* responses were concentrated in an early up-then-plateau trajectory (UF, 225 orthogroups), whereas *W. australiana* responses were concentrated in delayed trajectories that changed only by Day 4 (FU 655, FD 238, DF 245). Only ∼3.8% of orthogroups responsive in at least one species shared identical trajectories between the two species. These distributions indicate fundamentally different response tempos: *S. polyrhiza* mounts an immediate transcriptional response, reaching most of its final expression state within 3 h and maintaining it thereafter, whereas *W. australiana* barely reacts at 3 h and instead executes a slower, day-scale reorganization in which the bulk of transcriptional change accumulates over several days of continued exposure.

**Fig. 5.**
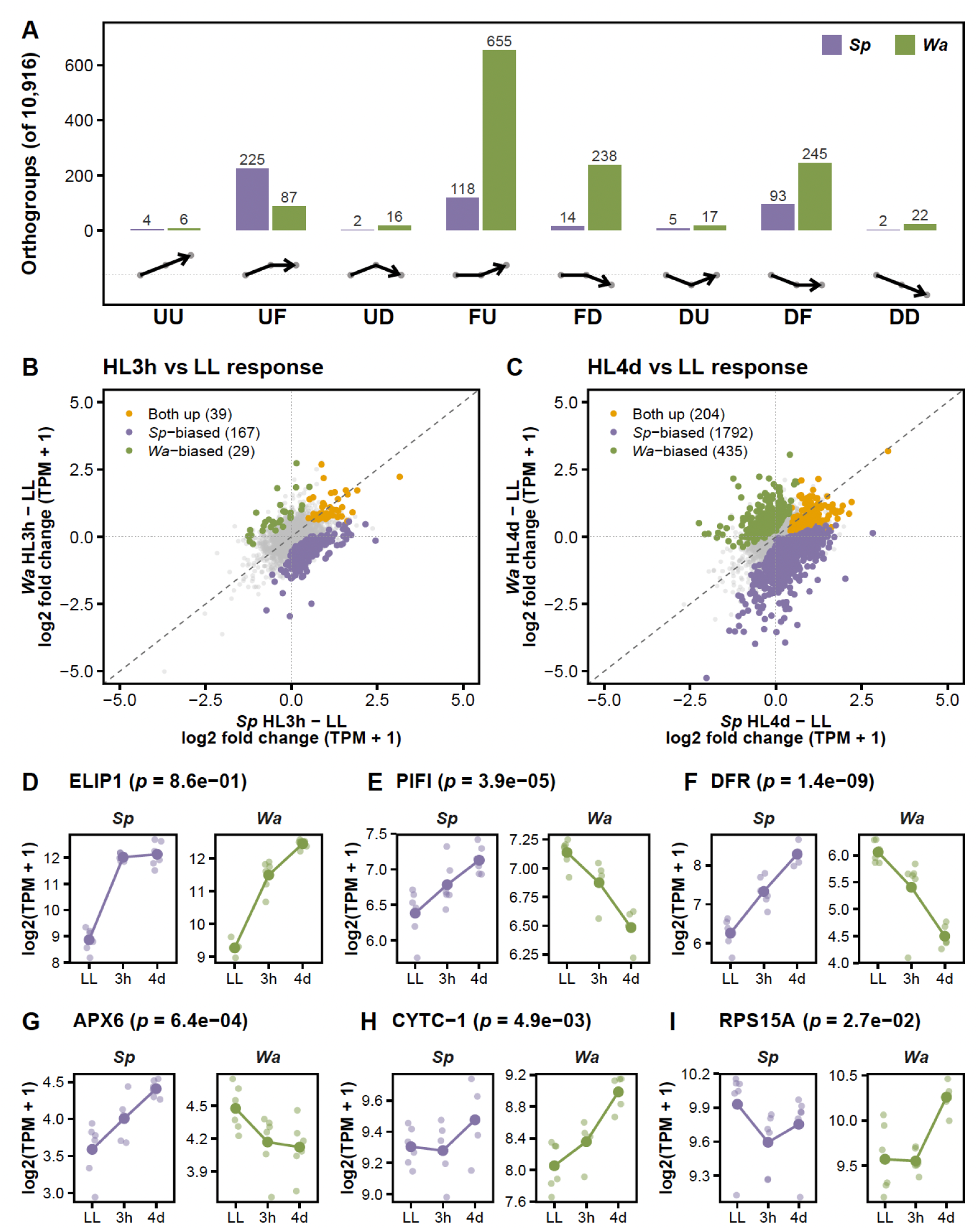
Transcriptomic responses to high light in *Spirodela polyrhiza* and *Wolffia australiana*. (A) High-light response trajectory frequency among 10,916 orthogroups expressed in both species. Each of the two consecutive transitions (LL→HL_3h and HL_3h→HL_4d) was independently classified as up (U), flat (F), or down (D) by within-species *limma* (BH-adjusted *p* < 0.05 and |log2 fold change| ≥ 0.5), yielding nine trajectory patterns. The FF (no-response) class, which dominates both species (*S. polyrhiza* 95.8%, *W. australiana* 88.2%), is not shown here; icon glyphs beneath each bar indicate the expected shape of each pattern (dashed line: baseline). Bars show the number of orthogroups per pattern in each species. Full lists are shown in Data S8-1. (B, C) Orthogroup response amplitude in *S. polyrhiza* (x) versus *W. australiana* (y) at HL_3h vs LL (B) and HL_4d vs LL (C). Each point is one orthogroup, colored by species × condition ANOVA (Benjamini–Hochberg-adjusted *p* < 0.05): orange, induced in both species (within-species *limma p* < 0.05 and log2 fold change > 0 in each); purple, *S. polyrhiza*-biased; green, *W. australiana*-biased; grey, not significant. Counts of each category are shown in the legend. Dashed line: 1:1 diagonal. (D–I) Representative gene expression profiles across LL, HL_3h (3h), and HL_4d (4d). Each panel splits *S. polyrhiza* (Sp, purple) and *W. australiana* (Wa, green). Lines connect within-species means; points show individual biological replicates (n = 6). The *p*-value in each panel title is the Benjamini–Hochberg-adjusted species × condition ANOVA interaction (HL_4d vs LL contrast). (D) ELIP1 (early light-induced protein), induced in both species. (E–G) *S. polyrhiza*-biased: PIFI (post-illumination chlorophyll-fluorescence increase; non-photochemical quenching), DFR (dihydroflavonol 4-reductase; anthocyanin biosynthesis), APX6 (ascorbate peroxidase 6; ROS scavenging). (H, I) *W. australiana*-biased: CYTC-1 (cytochrome *c*, mitochondrial Complex III→IV electron carrier), RPS15A (cytosolic small ribosomal subunit protein). Full GO enrichment results are provided in Data S8-2.

Among the 10,916 orthogroups, 204 were induced by HL_4d in both species (Fig. 5C; Data S8-1). GO enrichment of these shared high-light-responsive orthogroups highlighted carotenoid biosynthesis (GO:0016117; fold enrichment [FE] 17.6, p = 1.8 × 10⁻⁵; e.g., PSY1, ZDS1, ORLIKE), flavonoid biosynthesis (GO:0009813; FE 10.0, p = 3.3 × 10⁻⁴), the plastoglobule (GO:0010287; FE 13.9, p = 2.0 × 10⁻⁹), and the photosynthetic apparatus, including the chloroplast thylakoid membrane (GO:0009535; FE 6.5, p = 3.9 × 10⁻¹⁵; e.g., ELIP1), photosystem I assembly (GO:0048564; FE 8.3, *p* = 5.3 × 10⁻³), and the photosynthetic electron transport chain (GO:0009767; FE 6.1, p = 3.9 × 10⁻²) (Fig. 5D; Fig. S4; Data S8-2), indicating a shared core response to high light. Species-specific responses, however, differed markedly in both timing and function.

Consistent with its immediate tempo, *S. polyrhiza* had already engaged in a photoprotective, antioxidant program by HL_3h (Fig. 5A). At HL_3h, 167 orthogroups were Sp-biased versus only 29 Wa-biased (Fig. 5B); GO enrichment of these Sp-biased orthogroups highlighted responses to UV-B (GO:0010224; fold enrichment [FE] 8.2, p = 8.8 × 10⁻⁵), red light (GO:0010114; FE 9.5, p = 1.3 × 10⁻⁴), plastoglobule (GO:0010287; FE 6.9, p = 2.7 × 10⁻³), and L-ascorbate biosynthesis (GO:0019853; FE 16.7, p = 6.0 × 10⁻³; substrate for APX6) (Data S8-2). By HL_4d, the program expanded to 1,792 Sp-biased orthogroups (Fig. 5C, purple), enriched for the plastoglobule (GO:0010287), photosynthetic acclimation (GO:0009643), light-quality signaling (GO:0009646, GO:0071483), non-photochemical quenching (GO:0010196; e.g., PIFI, SOQ1, FLAP1), anthocyanin metabolism (GO:0046283; e.g., DFR, TT8), and reactive oxygen species metabolism (GO:0072593; e.g., APX6, CAT2, GPX2) (Fig. 5E–G; Fig. S5, Fig. S7; Data S8-1, Data S8-2).

In contrast, *W. australiana* barely responded at HL_3h: only 29 orthogroups were Wa-biased, and the sole enriched GO term was response to cytokinin (GO:0009735; FE 16.7, *p* = 3.3 × 10⁻³; e.g., YUC3, YUC8, ZFP8) (Fig. 5B; Data S8-2), matching the near-absent early trajectory response. By HL_4d, the Wa-biased cohort grew more than tenfold to 435 orthogroups (Fig. 5C, green), co-enriched for two coordinated biosynthetic modules. The mitochondrial respiratory chain was the single most significantly enriched, spanning Complex I (GO:0045271; FE 8.0, *p* = 5.3 × 10⁻⁹; e.g., FRO1/NDUFS4 and several unnamed subunits), Complex III (GO:0045275; FE 16.1, *p* = 6.7 × 10⁻⁶; e.g., AT5G25450, ATP6), and the mitochondrial intermembrane space (GO:0005758; FE 8.8, *p* = 4.0 × 10⁻³; e.g., CYTC-1). Co-enriched in parallel was the cytosolic ribosome and translational machinery (cytosolic large ribosomal subunit, GO:0022625; FE 6.0, *p* = 5.7 × 10⁻⁶; structural ribosomal constituent, GO:0003735; FE 3.3, *p* = 9.7 × 10⁻⁶; cytosolic small ribosomal subunit, GO:0022627; FE 5.2, *p* = 9.1 × 10⁻⁴; e.g., RPS15A) (Fig. 5H, I; Fig. S6; Data S8-2). When we further restricted the analysis to Wolffia’s delayed-up trajectory (FU pattern in Fig. 5A, 655 orthogroups), this ribosome enrichment became still more extreme (GO:0022625, GO:0022627, GO:0003735; all *p* < 10⁻³⁰; Data S8-2), indicating that ribosome biogenesis is the most tightly temporally coordinated component of *W. australiana*’s delayed response. The co-enriched photorespiration GO term (GO:0009853; FE 5.6, *p* = 1.8 × 10⁻³) comprised these Complex I (NADH dehydrogenase) subunits, annotated as photorespiratory through their role in NAD⁺ regeneration rather than as canonical photorespiratory enzymes. The co-enrichment of cytosolic ribosome biogenesis with mitochondrial oxidative phosphorylation, the biosynthetic and energetic core of proliferating cells, suggests that *W. australiana* prepares for accelerated growth by expanding translational output alongside the respiratory ATP supply that sustains it, rather than by activating photoprotective machinery.

### Lineage-specific metabolite accumulation in response to high light

To examine whether the contrasting high-light responses of *S. polyrhiza* and *W. australiana* were reflected at the metabolic level, we compared Day 4 metabolite profiles under LL and HL using LC-MS/MS, GC-MS/MS, and CE-TOF MS (Fig. 6; Fig. S8). These samples correspond to the time point used for the Day-4 HL transcriptome comparison, allowing changes in metabolite accumulation to be interpreted alongside gene expression responses. For each metabolite, we first calculated the HL-vs-LL log2FC within each species and tested whether the magnitude of this HL-associated response differed between species. We then examined whether these HL-associated responses corresponded to significant increases under HL within each species, using species-wise HL > LL tests after FDR correction (Data S9-1, S9-2).

**Fig. 6.**
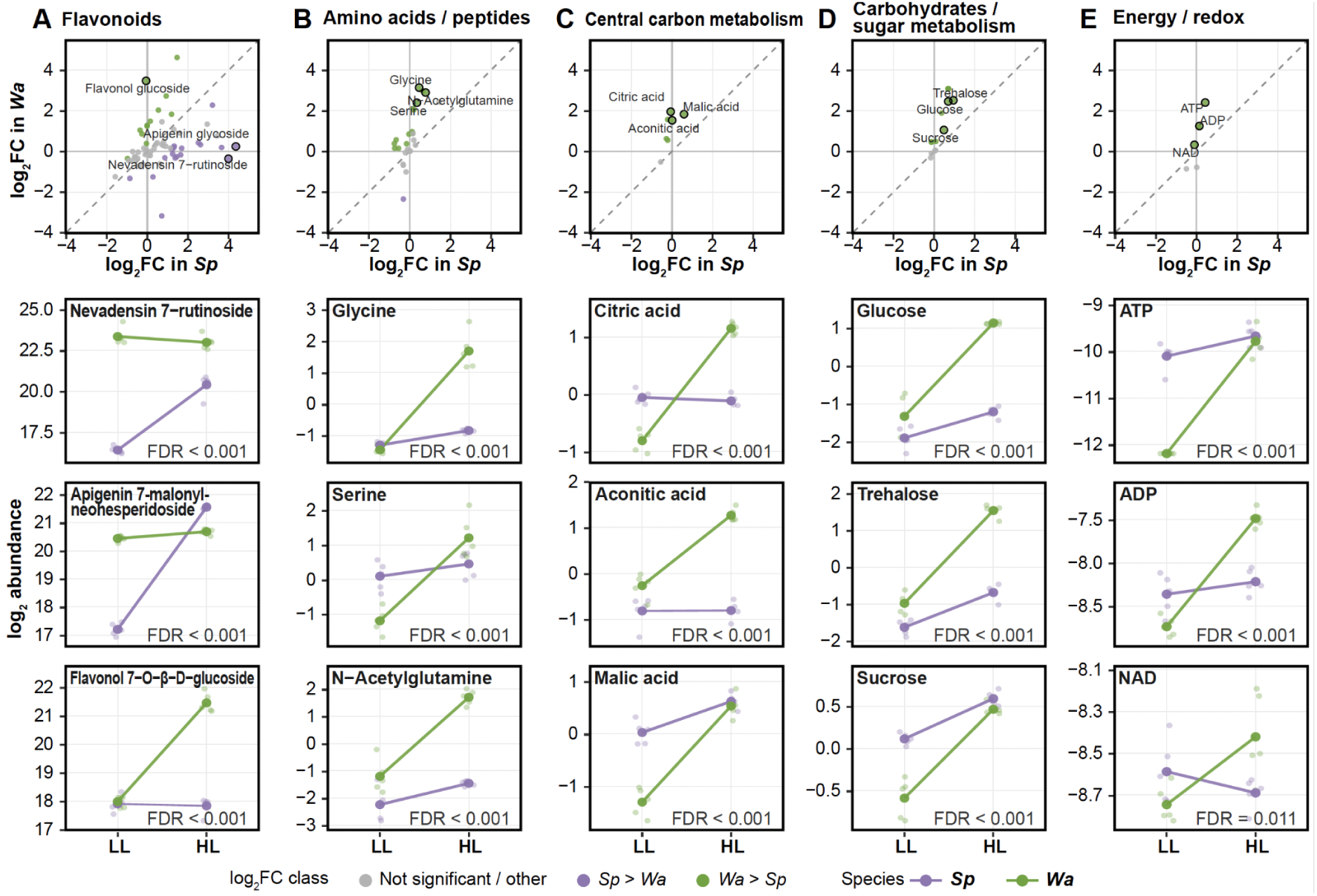
Species-specific metabolomic responses to high light in *Spirodela polyrhiza* and *Wolffia australiana*. (A-E) Category-level metabolomic responses to high light in flavonoids (A), amino acids/peptides (B), central carbon metabolism (C), carbohydrates/sugar metabolism (D), and energy/redox metabolites (E). In each panel, the left scatter plot shows the high-light log2 fold change for each metabolite in *S. polyrhiza* and *W. australiana*. Purple points indicate metabolites with a significant species x light interaction and a larger high-light response in *S. polyrhiza*, whereas green points indicate metabolites with a larger response in *W. australiana*. Gray points indicate metabolites without a significant species-biased high-light response. Black rings and labels identify the three representative metabolites shown in the abundance plots on the right. Right-side plots show log2 abundance under LL and HL for the representative metabolites; points indicate biological replicates and lines connect lineage means. FDR values indicate Benjamini-Hochberg-adjusted species x light interaction tests.

We first focused on LC-MS/MS flavonoid-related metabolites, because flavonoid biosynthetic processes were enriched in both species in the transcriptome analysis. Among the 61 flavonoid-related metabolites shown in Fig. 6A, the species-comparison analysis identified 18 metabolites with significantly larger HL-associated responses in *S. polyrhiza* and 12 with significantly larger responses in *W. australiana* after FDR correction (Fig. 6A; Data S9-2). We next investigated whether these species-biased HL-associated responses corresponded to significant HL-induced increases within each species. Six flavonoid-related metabolites were significantly increased under HL in both species, and all six were assigned to flavonol/flavone-related compounds, including apigenin-, quercetin-, kaempferol-, myricetin-, and luteolin-related annotations. Furthermore, among the 18 flavonoid-related metabolites with significantly larger HL-associated responses in *S. polyrhiza*, 16 were significantly increased under HL in *S. polyrhiza*. Similarly, among the 12 metabolites with significantly larger HL-associated responses in *W. australiana*, 11 were significantly increased under HL in *W. australiana* (Data S9-2). These results indicate that most species-biased HL-associated responses corresponded to actual HL-induced increases in the same species.

We next examined metabolite categories related to primary metabolism and cellular energy/redox status. The species-comparison analysis showed that several primary metabolic classes had larger HL-associated responses in *W. australiana* than in *S. polyrhiza* (Fig. 6B-E; Data S9-2). In amino acid- and peptide-related metabolites detected by GC-MS/MS, 14 metabolites showed significantly larger HL-associated responses in *W. australiana*, and 10 of these were significantly increased under HL in *W. australiana*; one metabolite showed a significantly larger HL-associated response in *S. polyrhiza*, and this metabolite was also significantly increased under HL in *S. polyrhiza* (Fig. 6B). Central carbon metabolism-related metabolites showed a strongly *W. australiana*-biased pattern: six metabolites had significantly larger HL-associated responses in *W. australiana*, and all six were significantly increased under HL in *W. australiana*, whereas no metabolite showed a significantly larger HL-associated response in *S. polyrhiza* (Fig. 6C). Sugar- and carbohydrate-related metabolites also showed stronger HL-associated responses in *W. australiana*: 10 metabolites showed significantly larger responses in *W. australiana*, and all 10 were significantly increased under HL in *W. australiana*, whereas no metabolite showed a significantly larger HL-associated response in *S. polyrhiza* (Fig. 6D). Energy/redox-related metabolites detected by CE-TOF MS included four metabolites with significantly larger HL-associated responses in *W. australiana*, and all four were significantly increased under HL in *W. australiana*, whereas no metabolite showed a significantly larger HL-associated response in *S. polyrhiza* (Fig. 6E). These results indicate that the primary metabolic categories showing species-biased HL-associated responses were strongly biased toward *W. australiana*, and that most of these *W. australiana*-biased responses corresponded to actual HL-induced increases in *W. australiana*.

Representative abundance plots further illustrated these lineage-biased metabolic responses (right three panels in Fig. 6A-E). Among flavonoids, *S. polyrhiza* showed stronger HL-associated increases in nevadensin 7-rutinoside and an apigenin derivative, whereas *W. australiana* showed a larger increase in a flavonol glucoside, indicating that flavonoid metabolism was also responsive in this species (Fig. 6A). In primary metabolism, glycine and serine showed stronger HL-associated increases in *W. australiana* (Fig. 6B). Because these metabolites are associated with photorespiration and one-carbon metabolism, this pattern may reflect differences in amino-acid-related metabolic adjustment under HL. *W. australiana*-biased increases were also observed for TCA-related organic acids, including citric acid, aconitic acid, and malic acid, as well as carbohydrates such as glucose, trehalose, and sucrose (Fig. 6C, D). In addition, ATP, ADP, and NAD showed larger HL-associated changes in *W. australiana*, consistent with the transcriptomic induction of mitochondrial oxidative phosphorylation genes (Fig. 6E). Overall, these results suggest that high-light acclimation in *S. polyrhiza* is characterized by a stronger flavonoid-associated response, whereas *W. australiana* shows broader remodeling of primary metabolism, carbohydrates, and energy/redox-related metabolites.

## Discussions

In the present study, we integrated phylogenomic, ecological, physiological, transcriptomic, and metabolomic analyses to explore the ecophysiological factors potentially associated with lineage-specific variation in molecular evolutionary rates in duckweeds. We confirmed substantial heterogeneity in branch lengths among duckweed lineages, suggesting lineage-specific variation in molecular evolutionary rate. We also found that lineages with and without prominent anthocyanin accumulation show different geographic distributions which are associated with molecular evolutionary rates. Anthocyanin-accumulating lineages were distributed more broadly, extending to higher latitudes and inland regions, whereas lineages without prominent anthocyanin accumulation were relatively more common in low- to mid-latitude regions. Furthermore, we revealed the difference in high-light acclimation strategies between anthocyanin-accumulation or non-accumulation lineages. Under HL conditions, anthocyanin-non-accumulating lineages maintained higher relative growth rates and higher PSII efficiency than anthocyanin-accumulating lineages. Detailed comparison between *Spirodela polyrhiza* and *Wolffia australiana*, representing lineages with and without prominent anthocyanin accumulation, respectively, further revealed contrasting acclimation strategies: *S. polyrhiza* preferentially activated photoprotective responses, including NPQ- and flavonoid-related pathways, whereas *W. australiana* maintained higher photochemical activity and showed stronger induction of mitochondrial oxidative phosphorylation and broader remodeling of primary metabolism. These results suggest that duckweed lineages have different high light acclimation strategies in how they balance light-energy dissipation, photochemical energy use, and downstream metabolic capacity. Taken together, we discuss the possibility that such ecophysiological divergence may be linked to lineage-specific molecular evolutionary rate variation.

### Divergent high-light acclimation strategies in duckweed lineages

Our physiological analyses suggested divergent high-light acclimation strategies in duckweed lineages. Under experimental HL conditions, anthocyanin-non-accumulating lineages showed higher relative growth rates and maintained higher Fv/Fm values than anthocyanin-accumulating lineages. Fv/Fm is widely used as an indicator of PSII maximum quantum efficiency and photoinhibitory damage stress (Maxwell and Johnson, 2000; Murchie and Lawson, 2013). In particular, *W. australiana* retained high PSII maximum efficiency after transfer to HL and showed strong recovery by Day 4. These results indicate that anthocyanin-non-accumulating duckweeds have evolved alternative physiological strategies that allow them to maintain photosynthetic performance and rapid growth under high-light conditions.

The detailed comparison between *S. polyrhiza* and *W. australiana* further revealed the differences between their acclimation strategies. In PAM analyses of plants grown under LL or HL conditions, *S. polyrhiza* showed greater regulated energy dissipation under high actinic light. Consistently, transcriptomic analyses showed that *S. polyrhiza* preferentially induced genes associated with oxidative-stress responses, NPQ, and anthocyanin/flavonoid biosynthesis. NPQ is a major photoprotective mechanism that dissipates excess light energy as heat and thereby reduces the photoinhibition under high light intensity (Niyogi, 1999; Ruban, 2016). Our metabolomic analyses also showed that HL-responsive flavonoids in *S. polyrhiza* included a dihydroflavonol intermediate and flavone-related derivatives, consistent with pigment-associated and antioxidant photoprotection. Therefore, *S. polyrhiza* appears to respond to high light primarily by enhancing photoprotective mechanisms that reduce excess light energy.

In contrast, PAM light-response analyses showed that *W. australiana* maintained higher Y(II) and ETR than *S. polyrhiza*, indicating that a larger fraction of absorbed light energy was used for PSII photochemistry. However, our RNA-seq analyses have not detected the transcriptional induction of photosynthesis-related genes in *W. australiana*. Instead, they showed stronger induction of two modules that were co-enriched in the same set of orthogroups: mitochondrial respiratory-chain and oxidative phosphorylation genes, including genes associated with Complex I, Complex III, and cytochrome c-mediated electron transport, and the cytosolic ribosome and translational machinery. Mitochondrial respiration is known to interact closely with photosynthesis by contributing to energy balance, redox homeostasis, and the dissipation of excess reducing equivalents under illuminated leaves (Raghavendra and Padmasree, 2003; Noguchi and Yoshida, 2008; Tcherkez et al., 2017). In our metabolomic analysis, *W. australiana* showed stronger HL-associated changes in amino-acid-related metabolites, TCA-related organic acids, sugars, sugar phosphates, and energy/redox-related metabolites such as ATP, ADP, and NAD. Together, the PAM, transcriptomic, and metabolomic analyses suggested that *W. australiana* maintains high photochemical activity by coupling PSII electron transport to mitochondrial energy metabolism and downstream primary metabolic sinks, rather than primarily dissipating excess light through pigment-based or NPQ-based photoprotection as seen in *S. polyrhiza*.

The two species also differed in the timing of their transcriptional responses. *S. polyrhiza* responded rapidly, with most expression changes occurring within 3 h, whereas *W. australiana* showed more changes by Day 4 (Fig. 5A). At 3 h, 167 orthogroups were already *S. polyrhiza*-biased and enriched for photoprotective and antioxidant functions, while only 29 were *W. australiana*-biased (Fig. 5B). This difference was also consistent with the fluorescence data. *W. australiana* maintained the highest Fv/Fm at Day 1 despite showing little transcriptional response at 3 h, suggesting that its early tolerance may rely on constitutive or post-transcriptional mechanisms. In contrast, *S. polyrhiza* showed a rapid transcriptional response but the lowest Fv/Fm at Day 1. Thus, the delayed response in *W. australiana* may reflect longer-term acclimation rather than an acute stress response.

### Mitochondrial respiratory and translational capacity may support high-light growth in *Wolffia*

The stronger response of oxidative phosphorylation and primary metabolism in *W. australiana* provided a possible explanation for how the species maintains high growth under HL conditions. When absorbed light exceeds the capacity for carbon fixation, plants need to prevent over-reduction of the photosynthetic electron transport chain. This process is achieved not only by dissipating excess energy through NPQ and photoprotective pigments, but also by increasing alternative sinks for reducing power and fixed carbon, including photorespiration, mitochondrial respiration, and carbohydrate metabolism (Ort and Baker, 2002; Bauwe et al., 2010; Noguchi and Yoshida, 2008; Ruban, 2016). Therefore, our data suggested that *W. australiana* uses such downstream metabolic capacity more strongly than *S. polyrhiza*. The Wa-biased induction of oxidative phosphorylation genes under HL-conditions suggest enhanced mitochondrial respiratory capacity, and the accumulation of ATP, ADP, and NAD is consistent with altered energy and redox metabolism. Furthermore, the stronger increases in accumulation of glycine and serine also suggest that amino-acid metabolism linked to photorespiration and one-carbon metabolism is more strongly affected by HL in *W. australiana*. Photorespiration is known to have a function as an important auxiliary pathway that consumes ATP and reducing power, maintains redox balance, and protects photosynthesis under conditions where CO₂ assimilation is limited relative to absorbed light energy (Ort and Baker, 2002; Bauwe et al., 2010). Similarly, the increases in citrate, isocitrate, aconitate, malate, glucose, trehalose, sucrose, and glucose 6-phosphate suggest broad remodeling of central carbon and carbohydrate metabolism. Because sink strength and carbohydrate partitioning are tightly linked to photosynthetic capacity and plant growth, these metabolic changes may support sustained carbon assimilation and biomass production under high light (Paul and Foyer, 2001; Smith and Stitt, 2007; Lemoine et al., 2013).

Furthermore, the parallel enrichment of cytosolic ribosome and translation-related genes suggests that this increase in respiration may have a role beyond simply dissipating excess energy. Ribosome biogenesis requires a large amount of energy and is closely related to growth rate (Sáez-Vásquez and Delseny, 2019). Therefore, the simultaneous induction of oxidative phosphorylation and ribosomal components may reflect increased biosynthetic activity rather than simply the disposal of excess reducing power. This interpretation is also supported by the timing of the response. When we focused on orthogroups that changed only between 3 hours and 4 days after transfer from low light to high light, ribosome enrichment became much stronger than in the Wa-biased cohort as a whole, suggesting that increased translational capacity is an important part of the delayed response in *W. australiana*. In this case, mitochondrial respiration in *W. australiana* may act not only as a sink for excess electrons, but also as an ATP source that supports increased biosynthetic activity.

This interpretation also explains an apparent discrepancy between transcriptomic and physiological or metabolomic results. Although photosynthesis-related genes were not broadly induced in *W. australiana* under HL conditions, they maintained higher Y(II) and ETR than *S. polyrhiza*. Transcriptomic profile does not always directly reflect photosynthetic capacity, because photosynthetic performance is known to be influenced by post-transcriptional regulation, electron transport regulation, and metabolic feedbacks (Pfannschmidt et al., 2009; Foyer et al., 2012). Thus, high-light acclimation in *W. australiana* may depend less on transcriptional upregulation of photosynthetic machinery and more on post-transcriptional, translational, post-translational, or metabolic regulation. The Wa-biased induction of ribosomal and translational components provides transcriptional evidence for this possibility, indicating that *W. australiana* expands translational capacity even where transcripts encoding the photosynthetic apparatus themselves are not induced. Such regulation could allow *W. australiana* to rapid adjustments of electron transport, respiratory capacity, and carbon utilization without large transcriptional changes in photosynthesis-related genes. Direct measurements of CO₂ assimilation, respiration, ROS production, and isotope-labeled carbon flux will be necessary to test this model.

### Linking high-light physiology to molecular evolutionary rate variation

The contrasting high-light responses observed in duckweeds provide a physiological explanation for understanding lineage-specific molecular evolutionary rate variation. Previous studies have demonstrated that molecular evolutionary rates are associated with life-history traits such as growth form, generation time, and plant height in flowering plants (Smith and Donoghue, 2008; Lanfear et al., 2013; Smith et al, 2026). These studies suggest that molecular rate variation in plants may partly reflect differences in growth-associated processes, rather than being determined solely by generation time or population-genetic factors. This perspective is especially relevant to duckweeds because they reproduce clonally through repeated budding of fronds and show exceptionally rapid growth speed (Landolt, 1986; Sree et al., 2015; Ziegler et al., 2015). In our phylogenomic analyses, the duckweed lineages showed substantial variation in branch length and the three lineages that do not accumulate anthocyanins, *Wolffiella hyalina*, *L. aequinoctialis*, and *W. australiana*, occupied the three highest positions, with all six lineages significantly different from one another (Fig. 1C) Notably, the *Wolffia/Wolffiella* clade maintained higher growth rates under HL conditions. Because plants are generally considered not to segregate the germline early in development, unlike many animals, somatic mutations during vegetative growth could potentially contribute to heritable genetic variation (Lanfear, 2018; Schoen and Schultz, 2019). Therefore, sustained growth under high light could increase mutations associated with DNA replication, providing a link between high-light physiology and molecular evolutionary acceleration.

This association is unlikely to be explained simply by phylogenetic position. The anthocyanin-non-accumulating lineages do not form a single clade, as *L. aequinoctialis* is sister to the anthocyanin-accumulating *L. gibba* (Fig. 1B), indicating at least two independent changes in anthocyanin status. This sister pair also shows the same pattern: *L. aequinoctialis* had a higher relative evolutionary rate than *L. gibba* (Fig. 1C). A similar pattern was also observed for geographic distribution within Lemna (Fig. S1). Thus, the rate difference appears to be associated with anthocyanin status rather than with a particular phylogenetic position.

In addition to growth-associated DNA replication, redox-dependent processes under high-light environments may also influence the accumulation of mutations. Excess light can over-reduce the photosynthetic electron transport chain and promote the generation of reactive oxygen species (ROS), which damage cellular components, including DNA (Foyer and Shigeoka, 2011; Khorobrykh et al., 2020; Roldán-Arjona and Ariza, 2009). Our results suggested that *W. australiana* and other lineages without prominent anthocyanin accumulation may mitigate high-light stress through active photosynthetic and respiratory metabolism. In these lineages, higher photochemical activity, stronger oxidative phosphorylation responses, and broader changes in primary metabolism may help maintain electron flow and energy balance under high-light environments, thereby reducing the accumulation of excess excitation energy. At the same time, sustained high photochemical activity could increase ROS production. Thus, the differences in high-light acclimation among duckweeds lineages could contribute to variation of molecular evolutionary rates through two non-mutually exclusive pathways: a replication-mediated pathway associated with rapid vegetative growth, and a redox-mediated pathway associated with photosynthetic electron transport, mitochondrial respiration, oxidative stress, and DNA damage/repair balance.

Importantly, our study does not directly measure mutation rates, and therefore we cannot determine whether the observed branch-length differences are caused by higher mutation rates associated with differences in high-light response strategies. Our data do, however, argue against variation in selective constraint as the main explanation. Median *d_N_*scaled proportionally with median *d_S_* across all six lineages (Fig. 1E), and the ranking of lineages by ω did not follow their ranking by evolutionary rate: the slowest-evolving lineage, *S. polyrhiza*, had the highest ω of the six (0.56), the fastest-evolving lineage, *Wolffiella hyalina*, had the second highest (0.53), and the lowest value was found in the intermediate-rate *L. gibba* (0.43). The several-fold rate differences among duckweed lineages are therefore more consistent with variation in mutation rate than with differences in purifying selection. Our sampling includes only six lineages and two inferred changes in anthocyanin status, limiting the number of independent comparisons. Other traits that changed alongside anthocyanin loss, including body size, frond structure, root loss, and clonal growth rate, therefore cannot be disentangled from it. Denser sampling within *Lemna*, where anthocyanin status varies at shallower phylogenetic depths, will be needed to separate these effects. According to the neutral and nearly neutral theories, substitution rates are shaped by mutation rates, natural selection, and effective population size; thus, variation in molecular evolutionary rates can arise through multiple population-genetic mechanisms (Kimura, 1983; Ohta, 1992). Our data therefore constrain the selective component but cannot exclude a contribution from differences in effective population size. The inferred link between high-light acclimation and molecular evolutionary acceleration should therefore be interpreted as a hypothesis-generating result rather than direct evidence of causality. Nevertheless, the present study provides experimental evidence for concordant patterns among phylogenomic, physiological, transcriptomic, and metabolomic analyses, supporting the idea that ecological and physiological differences contribute to lineage-specific molecular rate heterogeneity in plants.

The evolution of duckweed lineages involved drastic reductions in body size and structural complexity, culminating in *Wolffia* lineage, the smallest flowering plants of the world (Landolt, 1986; Nauheimer et al., 2012; Sree et al., 2015). Our results suggested the changes in high-light acclimation strategy in addition to this morphological simplification. Early-diverging lineages with anthocyanin accumulation rely mainly on pigment-based photoprotection, whereas derived anthocyanin-non-accumulating lineages appear to maintain photosynthetic activity and use light-derived energy in their growth and primary metabolism. This strategy against high light may facilitate the adaptation to open aquatic habitats and the rapid clonal expansion.

More broadly, our findings provided insights that molecular evolutionary rate variation in plants may be linked not only to life-history traits, such as generation time and plant size, but also to physiological mechanisms of photosynthesis to use and process environmental light energy. In duckweeds, the strategies to sustain photochemical activity and rapid growth under high light may increase metabolic turnover accompanied by mutation accumulation. Similar links between environmental physiology and molecular evolutionary acceleration have been proposed in other aquatic or highly specialized plant lineages, including Podostemaceae and Lentibulariaceae (Ibarra-Laclette et al. 2011; Katayama et al., 2022). Future studies under controlled light environments, combined with direct measurements of somatic mutations, DNA damage, photosynthetic carbon assimilation, respiration, ROS dynamics, and metabolic flux, will be necessary to test whether high-light physiology causally influences mutation rates and molecular evolutionary rates in duckweeds.

## Resource availability

## Lead contact

Supplemental information is available for this paper. Correspondence and requests for materials should be addressed to the lead contact, Natsu Katayama.

## Materials availability

Duckweed accessions used in this study are listed in Supplementary Data S0-1. Materials obtained from public stock collections should be requested from the corresponding collection when available. Other materials may be made available from the lead contact upon reasonable request, subject to availability and institutional or legal requirements.

## Data and code availability

The RNA-seq raw reads and transcriptome assemblies have been deposited in the NCBI under BioProject PRJNA1508391 (Reviewer link: https://dataview.ncbi.nlm.nih.gov/object/PRJNA1508391?reviewer=gn48ps7d1cnfbhv8919s56a7r6).

## Acknowledgments

We are grateful to Dr. T. Oyama and Dr. S. Ito at Kyoto University and Dr. T. Muranaka at Nagoya University for providing duckweed accessions. We also thank Dr. S. Kuraku for support with computational resources. The computations were performed using the computer facilities of the National Institute for Basic Biology and the Research Center for Computational Science (Project: 25-IMS-C320). This work was supported by JSPS KAKENHI Grant Numbers 22KJ0490 to N.K. and 24K23197 to Y.W.K., the Naito Foundation Research Grant for Female Researchers to N.K., and the National Institute of Genetics (NIG) Postdoctoral Research Fellowship to Y.W.K.

## Author contribution

Conceptualization, N.K., Y.W.K., and M.I.; Methodology, N.K., Y.W.K., M.K., K.K., M.Y.H., and M.I.; Investigation, Y.W.K., N.K., M.K., M.I., K.K., H.T., R.S., E.J., and L.V.; Formal analysis, Y.W.K. and N.K.; Metabolomic analysis, K.K., H.T., R.S., A.O., and M.Y.H.; Resources, M.I., and N.K.; Data curation, Y.W.K. and N.K.; Visualization, Y.W.K. and N.K.; Writing – original draft, N.K. and Y.W.K.; Writing – review & editing, all authors; Supervision, N.K.; Funding acquisition, N.K. and Y.W.K.

## Methods details

### Plant materials and growth conditions

Six duckweed species representing 5 genera of Lemnoideae were mainly used in this study: *Spirodela polyrhiza* “Sp7498” (Sp), *Landoltia punctata* “Lp9387” (Lp), *Lemna gibba* “Lgp8L” (Lg), *Lemna aequinoctialis* “M10E” (La), *Wolffiella hyalina* “Wh9525” (Wh), and *Wolffia australiana* “Wa8730” (Wa). Species and strain information are listed in Supplementary Data S0-1. Plants were maintained under axenic culture conditions in NF liquid medium with 1% sucrose under approximately 50µmol m⁻² s⁻¹ photosynthetic photon flux density (PPFD) at 25°C under a 16 h light / 8 h dark photoperiod before experiments.

For light-treatment experiments, plants were precultured under low-light conditions (LL; approximately 50–100 µmol m⁻² s⁻¹ PPFD) under a 16 h light / 8 h dark photoperiod. High-light treatment was performed by transferring plants to high-light conditions (HL; approximately 800–1000 µmol m⁻² s⁻¹ PPFD) under the same temperature and photoperiod conditions. Unless otherwise stated, physiological measurements and sampling were conducted using plants grown under LL and plants exposed to HL for the indicated duration.

### Classification of anthocyanin-accumulating and anthocyanin-non-accumulating lineages

Duckweed lineages were classified as anthocyanin-accumulating or anthocyanin-non-accumulating based on previously reported flavonoid profiles on Lemnoideae. In particular, we followed the presence/absence information for anthocyanins summarized by Les *et al*. (1997), which incorporated flavonoid data originally obtained by McClure and Alston (1966).

### Sequence data preparation

Protein and coding sequences (CDS) were obtained for the six representative duckweed species utilized in subsequent phylogenomic and molecular analyses: *S. polyrhiza*, *Landoltia punctata*, *L. aequinoctialis*, *L. gibba*, *W. australiana*, and *Wolffiella hyalina*.

For *S. polyrhiza*, a comprehensive mapping reference was generated by integrating the nuclear reference genome from the Lemna Genome Hub (Sp_polyrhiza_9509-REF-OXFORD-3.0; Ernst et al. 2025) with the corresponding plastid (NC_015891) and mitochondrial (NC_017840) genomes. Similarly, for *W. australiana*, the nuclear assembly (Wo_australiana_8730-REF-CSHL-1.0; Ernst et al. 2025) was supplemented with its chloroplast genome (NC_015899). Mitochondrial gene models for *W. australiana* were identified using MFAnnot under default parameters (https://github.com/BFL-lab/Mfannot). Organellar gene models for both species were retrieved from NCBI and incorporated into their respective combined genomic annotations.

For *L. gibba*, the chromosome-level reference assembly and gene annotation were obtained from the Lemna Genome Hub (Le_gibba_7742a; Ernst et al. 2025). All three genome-based annotations (*S. polyrhiza, W. australiana, L. gibba*) were standardized using AGAT agat_convert_sp_gxf2gxf.pl, and protein and CDS sequences were extracted using AGAT agat_sp_extract_sequences.pl, retaining one representative (longest) transcript per gene (Dainat et al., 2026).

For *Landoltia punctata*, *L. aequinoctialis*, and *Wolffiella hyalina*, protein and CDS sequences were inferred from *de novo* transcriptome assemblies. Raw paired-end RNA-seq data for all three species were generated in this study using strand-specific libraries sequenced on an Illumina NovaSeq 6000 platform (150 bp paired end) by Azenta Life Sciences (Tokyo, Japan) (Data S0-1). In each instance, transcripts were assembled via Trinity v2.15.2 with default settings (Grabherr et al., 2011; Haas et al., 2013), which include in-silico normalization of reads to a maximum coverage of 200, and clustered at 98% identity using CD-HIT-EST v4.8.1 to minimize redundancy (Fu et al., 2012). Coding regions were predicted using TransDecoder v5.5.0 (https://github.com/TransDecoder/TransDecoder); candidate open reading frames (ORFs) were validated against Pfam-A using HMMER v3.1b2 and UniProt/Swiss-Prot using DIAMOND v2.2.5 (Mistry et al., 2021; Eddy, 2011;The UniProt Consortium, 2023; Buchfink et al., 2021). Final predictions were generated using TransDecoder.Predict, retaining only ORFs supported by homology or Pfam domains, with the single representative isoform for each Trinity gene selected based on the highest coding score.

### Orthologous grouping across land plant species

Orthologous gene groups (OGs) were inferred using OrthoFinder v3.1.3 with the default parameters from protein sequences of eight plant species (Emms & Kelly, 2019; Emms et al., 2025): *Arabidopsis thaliana*, *Colocasia esculenta* (L.) Schott, *S. polyrhiza*, *Landoltia punctata*, *L. aequinoctialis*, *L. gibba*, *W. australiana*, and *Wolffiella hyalina*. For *A. thaliana*, the Araport11 representative gene models (one peptide per gene; release 2025-02-14) were downloaded from TAIR (Cheng et al., 2017). For *C. esculenta*, the gene model and the genome assembly were obtained from the China National GeneBank Database (Yin et al., 2021; accession CNP0001082). Sequences for the six Lemnoideae species were prepared as described above.

### Phylogenetic tree and branch length

Phylogenetic relationships and relative molecular evolutionary rates within Lemnoideae were inferred using six Lemnoideae species with *C. esculenta* as an outgroup (seven taxa in total). From the OrthoFinder output, we identified 2,819 single-copy orthogroups in which all seven species were represented by exactly one gene copy. For each orthogroup, protein sequences were aligned with MAFFT v7 (--inputorder; Katoh & Standley, 2013), and the resulting alignments were used to guide codon-based CDS alignments using Tranalign (EMBOSS; Rice et al., 2000). Gene trees were inferred using IQ-TREE 3 (TIM3+F+I; -B 1000; Minh et al., 2020) with *C. esculenta* as the outgroup. A species tree was also estimated from a concatenated supermatrix of all codon alignments, constructed using AMAS with a partition model per orthogroup (Borowiec, 2016), inferred using IQ-TREE 3 (-B 1000), and edited using PhyloWeaver (Kawaguchi Y.W, 2026)

To quantify branch lengths, root-to-tip distances were computed from each gene tree using a custom script based on ETE3 v3.1.3, with *C. esculenta* as the outgroup (Huerta-Cepas et al., 2016). The base node was defined as the internal node representing the split between *C. esculenta* and the six Lemnoideae species. Branch lengths were normalized by dividing each species’ base-node-to-tip distance by that of *C. esculenta* in the same tree, yielding a relative branch length that accounts for gene-specific rate variation. Differences in relative branch length among lineages were tested with the Friedman test (species as treatments, gene trees as blocks), followed by post-hoc pairwise Wilcoxon signed-rank tests with Holm correction, in R v4.4.1 (R Core Team, 2024).

### Estimation of dN/dS across lineages

To assess whether the among-lineage variation in evolutionary rate reflects differences in selection pressure, we estimated synonymous (*d_S_*) and non-synonymous (*d_N_*) substitution rates and the *d_N_*/*d_S_*ratio (ω) for each of the 2,819 single-copy orthogroups. Using the codon alignments and gene trees described above, we fitted the MG94×REV codon model to every branch with HyPhy v2.5.61 (Kosakovsky Pond et al., 2020), obtaining a per-branch baseline ω together with the synonymous and non-synonymous branch lengths; in-frame stop codons were masked prior to fitting. Orthogroups were retained for analysis only if, in all seven taxa, the baseline branch length was between 0.005 and 1.0 substitutions/site, the baseline ω was ≤ 2, *d_S_*was between 0.01 and 1.0, the codon alignment contained no in-frame stop codons and less than 80% gaps, and the alignment was at least 100 codons long; orthogroups inferred to be under selection on three or more of the eight branches, a signature of alignment error, were also removed. This retained 2,474 orthogroups. Because *d_N_*/*d_S_* estimated on a single branch is unreliable when synonymous divergence is low, comparisons among lineages were based on terminal branches, which are phylogenetically independent, and branches with an inferred selected-class ω greater than 5 were excluded to avoid inflation by episodic positive selection. For each lineage we summarised *d_N_* and *d_S_* as the median across orthogroups, and assessed proportionality (constant ω) by regression of median *d_N_*on median dS through the origin.

### Global occurrence data and distribution analysis

Occurrence records of duckweed species were obtained from GBIF (GBIF.org, 19 September 2024) for the five genera of Lemnoideae: *Spirodela*, *Landoltia*, *Lemna*, *Wolffiella*, and *Wolffia*. Records were assigned to species using the binomial species name extracted from the scientificName field, and only records assigned to accepted 36 duckweed species based on the updated Lemnoideae taxonomic key were retained (Bog et al., 2020a, 2020b). Occurrence records lacking species-level identification or geographic coordinates were removed. Records with latitude or longitude values outside valid geographic ranges, records located at 0° latitude and 0° longitude, and duplicate records with identical species name and coordinates were also excluded. To reduce the influence of poorly localized records, occurrences with coordinate uncertainty greater than 10 km were removed, while records without coordinate uncertainty information were retained. To reduce spatial sampling bias, the remaining records were spatially thinned by retaining one randomly selected occurrence per species within each 0.5° latitude x 0.5° longitude grid cell.

After filtering and spatial thinning, the dataset contained 18,227 occurrence records from 29 duckweed species. Global distribution maps were generated by plotting thinned occurrence records on a world map. All analyses and visualizations were performed in R using tidyverse, maps, viridis, and ggbeeswarm.

### Growth rate measurement under different light condition

To compare growth responses under different light conditions, plants were grown under LL and HL conditions for seven days. Frond area was recorded as digital photos every 24 h using six duckweed lineages (Sp, Lp, Lgp8L, M10E, Wh, and Wa) grown under low light and high light conditions (PFD100 and PFD1000). Frond area was extracted from cropped well images using OpenCV-based thresholding in 8-bit CIELAB color space. Candidate masks included a primary threshold of L = 0–255, a = 90–135, and b = 155–255, which was used to detect green fronds. Because this primary threshold occasionally underestimated frond area in images with high-light-induced yellowing or reddening, low brightness, or reflection-induced loss of internal frond pixels, broader candidate masks were also generated using L = 36–255, a = 0–138, and b = 143–255. For reddish high-light fronds, an additional red/orange threshold, L = 0–255, a = 136–255, and b = 134–244, was combined with the broader mask by logical OR. Candidate masks were optionally processed by removing small noise components, reflection artifacts, and thin root-like line structures. Low-brightness images were also evaluated after CLAHE-based brightness correction. The best-matching mask was selected manually for each image and the selected image-level frond areas were used for following analyses.

The Day 0–Day 4 interval was selected as the primary growth window because inspection of the Day 0–Day 7 growth trajectories indicated that this period captured the early growth phase before stronger late-stage deceleration or saturation in some wells. Relative growth rate (RGR) was calculated using the selected image-level frond areas as:

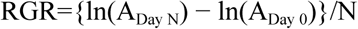

where A_Day 0_ and A_Day N_ represent total frond area at Day0 and DayN, respectively.

As a complementary analysis, model-based growth rates were estimated from the full Day 0–Day 7 time series for each well. Frond area trajectories were fitted separately for each well using exponential and logistic growth models. The exponential model was fitted as ln(area) = intercept + r × time, where r represents the exponential growth rate per day. The logistic model was fitted to the untransformed frond area trajectory to account for possible saturation at later time points. Exponential-fixed, logistic-fixed, and AIC-based model-selection strategies were evaluated. In the AIC-based strategy, the model with the lower AIC was selected for each well. These model-based growth-rate estimates were used as supplementary analyses to evaluate whether the main Day 0–Day 4 RGR results were robust to alternative growth-rate estimation approaches.

Both interval-based RGR and model-based growth rate estimates (r) were analyzed using Gamma generalized linear mixed models with a log link. For species-level analyses, the model included the fixed effects of lineage, light condition, and their interaction, with experimental batch as a random effect:

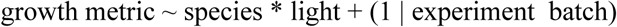

For anthocyanin-status analyses, lineages were grouped as anthocyanin-accumulating (Sp, Lp, Lgp8L) or non-anthocyanin (M10E, Wh, Wa), and the model included anthocyanin status, light condition, and their interaction as fixed effects, with lineage and batch as random effects:

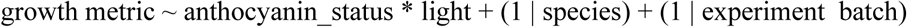

In these models, the response variable was either RGR or the fitted growth rate *r*. Models were fitted using lme4::glmer with a Gamma error distribution and log link (nAGQ = 0). Type II Wald chi-square tests were performed using car::Anova, and pairwise contrasts between anthocyanin-accumulating and non-anthocyanin groups within each light condition were estimated using emmeans on the response scale.

### Measurement of maximum quantum yield of PSII photochemistry

To evaluate photoinhibition under high light, the maximum quantum yield of PSII photochemistry (Fv/Fm) was measured in six representative species. Plants were precultured under LL conditions for 3 days and then transferred to HL conditions. Fv/Fm was measured at Day 0 to Day 4 after transfer to HL. Chlorophyll fluorescence was measured using a pulse-amplitude-modulated fluorometer in the dark period. Fv/Fm was calculated as:

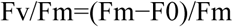

where F0 is minimum fluorescence and Fm is maximum fluorescence after a saturating light pulse.

Fv/Fm values at Day 0, Day 1, and Day 4 after transfer to HL were analyzed using a linear model including lineage, time point, and their interaction as fixed effects. Time point was treated as a categorical factor. Significance of fixed effects was evaluated using Type II ANOVA. Pairwise comparisons among accessions within each time point and among time points within each accession were performed using estimated marginal means with Tukey adjustment.

### Photosynthetic light-response and energy partitioning analyses

Photosynthetic light responses were compared between *S. polyrhiza* and *W*. *australiana* after plants had been grown under LL or HL conditions for four days. Chlorophyll fluorescence parameters were measured across a series of AL intensities using a pulse-amplitude-modulated fluorometer. Electron transport rate through PSII (ETR) was calculated from the effective quantum yield of PSII [Y(II)] and incident AL intensity as ‘ETR = Y(II) x PAR x 0.84 x 0.5’.

Energy partitioning of absorbed light was evaluated from the quantum yields Y(II), Y(NPQ), and Y(NO), which represent PSII photochemistry, regulated non-photochemical energy dissipation, and non-regulated energy dissipation, respectively. These parameters satisfy the relationship Y(II) + Y(NPQ) + Y(NO) = 1.

For quantitative comparisons of energy partitioning, two representative ALt intensities were extracted from the light-response measurements: PPFD = 61.885 µmol m⁻² s⁻¹ as the low-AL point and PPFD = 1012.500 µmol m⁻² s⁻¹ as the high-AL point. These measurement points are referred to as PPFD_50 and PPFD_1000, respectively. For each quantum-yield parameter, species differences were tested within each combination of growth light condition and AL intensity using linear mixed-effects models, followed by pairwise comparisons of estimated marginal means with Sidak adjustment.

### Gene expression analysis

We obtained transcript data for *S. polyrhiza* and *W. australiana* under three conditions (LL: control light condition, HL_3h: 3h after transfer from low light to high light, HL_4d: four days after transfer from low light to high light) with six biological replicates per condition. For each species and condition, three replicates were sequenced by Azenta Life Sciences (Tokyo, Japan) using strand-specific libraries on an Illumina NovaSeq 6000 platform (150 bp paired end), whereas the remaining three replicates were sequenced by BGI (Hong Kong, China) using strand-specific libraries on a DNBSEQ-G400 platform (150 bp paired end). After trimming using fastp v1.0.1 with the parameters --length_required 10 --trim_poly_x (Chen et al. 2018), these reads were mapped to species-specific mapping references using hisat2 v2.2.1 with the default parameter (Kim et al. 2019). These reference genomes were prepared as described above.

Read counts were obtained using featureCounts v2.0.8 with the parameters -p -t exon -g Parent -M --fraction -O, and were normalized to transcripts per million (TPM; Liao et al. 2014). With these options, multi-mapping reads (-M) and reads overlapping more than one transcript isoform of the same gene (-O) were retained but assigned fractionally (--fraction): each such read contributed a fractional count of 1/N distributed across its N alignments or overlapping isoforms, rather than being discarded as ambiguous or counted in full against every target. Because recently duplicated paralogs and exons shared among isoforms of the same gene generate many such reads, this fractional assignment avoids both the loss of these reads and their double-counting. Expression values were then summarized at the orthogroup level by summing TPM across paralogs within each orthogroup. The orthogroups used for expression summarization were defined using orthologous grouping across eight land plant species, including *A. thaliana* and other outgroup taxa, as described below. For statistical analyses, log-transformed expression values were computed as log2(TPM + 1) and corrected for batch effects using the removeBatchEffect function from the limma package v3.62.2 in R v4.4.1 (Ritchie et al. 2015; R Core Team, 2024). The model matrix included species (*S. polyrhiza* vs *W. australiana*) and treatment condition (LL, HL_3h, HL_4d), while sequencing batch (Illumina NovaSeq 6000 vs DNBSEQ-G400) was incorporated as a covariate to remove platform-specific technical variation.

To detect orthogroups exhibiting species-specific expression responses to high-light treatment, we performed two-way ANOVA for each orthogroup and tested the interaction between species and treatment condition. Planned contrasts included HL_4d vs HL_3h, HL_3h vs LL, and HL_4d vs LL. *p*-values were adjusted for multiple testing using the Benjamini-Hochberg procedure, and orthogroups with significant species–condition interactions were defined as differentially responsive orthogroups (FDR < 0.05).

### Gene Ontology (GO) enrichment analysis

The GO term of each OG was determined based on *Arabidopsis thaliana* (https://www.arabidopsis.org/download/list?dir=GO_and_PO_Annotations%2FGene_Ontology_Annotations, Access on 23 February 2026). The enrichment analysis was performed using topGO v.2.58.0 with the ‘weight01’ algorithm (Alexa et al., 2006), which takes into account the hierarchical structure of GO terms, along with GO.db version 3.20.0 (Carlson, 2019) in R v4.4.1. GO terms were considered enriched when the weight01 Fisher *p* value was below 0.05 and the term was annotated to at least 10 orthogroups in the tested universe, with at least 3 orthogroups in the query set. Fold enrichment (FE) is reported as the ratio of observed to expected orthogroup counts. Because fold enrichment scales inversely with the size of the query set, no fold-enrichment cutoff was applied. Complete results for all comparisons and cohorts, unfiltered, are provided in Data S8-2.

### Metabolome analysis

Metabolite profiling was performed for *S. polyrhiza* and *W. australiana* using Day 4 samples collected under LL and HL conditions. These samples corresponded to the time point used for the HL_4d RNA-seq analysis, enabling direct comparison between transcriptomic and metabolomic responses. Tissue samples were collected in microtest tubes, measured fresh weight (around 50 mg), and immediately frozen in liquid nitrogen. Untargeted LC–MS/MS analysis was performed by BGI Genomics Co., Ltd. For LC–MS/MS, metabolite extraction, peak extraction, metabolite identification, data preprocessing, and quality control were conducted by BGI Genomics. Metabolite extraction and subsequent analyses using GC–MS/MS and CE–TOF MS were conducted as previously described (Oikawa et al., 2011a; 2011b; 2023; Kawade et al., 2025), with five biological replicates per species and treatment. To avoid undefined values during log transformation, a metabolite-specific pseudocount was added to each metabolite, defined as half of the smallest positive value observed for that metabolite within the dataset. Metabolite abundances were then transformed as log2(value + pseudocount). For each metabolite, mean log2 abundance was calculated separately for each lineage and light treatment, and HL-associated changes were quantified within each species as log2FC = log2HL_mean − log2LL_mean. To identify species-specific metabolite responses to high light, we fitted a linear model for each metabolite in the form log2y ∼ Lineage * Light and tested the significance of the Lineage × Light interaction. P values were adjusted across metabolites using the Benjamini–Hochberg procedure, and metabolites with FDR < 0.05 were considered significantly differentially responsive between the two species. Significant metabolites were further classified according to whether HL-associated changes were greater in *W. australiana* than in *S. polyrhiza* (Wa > Sp) or greater in *S. polyrhiza* than in *W. australiana* (Sp > Wa). For interpretation and visualization, metabolites detected by each MS platform were assigned to functional categories based on metabolite identity and known pathway annotations. In the main comparisons shown in Figure 6, we focused on metabolite groups directly relevant to flavonoid metabolism, photorespiration, central carbon metabolism, and carbohydrate/sugar metabolism.

## Declaration of generative AI and AI-assisted technologies in the manuscript preparation process

During the preparation of this work, the authors used ChatGPT by OpenAI to assist with English editing, wording refinement, and improving manuscript clarity. The authors also used Codex by OpenAI and Claude Code by Anthropic to assist with drafting and debugging analysis scripts. All AI-assisted text and code were reviewed, edited, and verified by the authors, who take full responsibility for the content of the published article.

